# Phosphorylation modulates estrogen receptor disorder by altering long-range hydrophobic interactions

**DOI:** 10.1101/2023.07.14.548966

**Authors:** Zhanwen Du, Han Wang, Chen Wu, Matthias Buck, Wenwei Zheng, Alexandar L. Hansen, Hung-Ying Kao, Sichun Yang

**Author notes:** These authors contributed equally to this work (Zhanwen Du, Han Wang).

## Abstract

Protein intrinsic disorder is coupled to a range of biological phenomena, from gene regulation to cancer progression. Phosphorylation of the estrogen receptor (ER) at Ser118 through its disordered N-terminal domain (NTD) activates its transcriptional function, but it is challenging to rationalize how this modification regulates ER activity. Using biophysical approaches of small-angle X-ray scattering and nuclear magnetic resonance spectroscopy, we demonstrate that Ser118 phosphorylation triggers long-range conformational changes in ER-NTD, particularly between two hydrophobic clusters of residual structures. Alanine substitution of hydrophobic amino acids near Ser118 produces similar conformational alterations and rescues impaired ER activity caused by a phosphorylation-deficient mutant. These findings establish a direct link between phosphorylation-induced conformational changes and the activation function of this disordered protein as a promising avenue to block ER transcriptional activation.

## Introduction

Transcription factors, such as p53, androgen receptor (AR), and estrogen receptor (ER), are master regulators of gene expression^1,2^. The N-terminal domain (NTD) of these transcription factors has emerged as a promising target for cancer therapy due to its role in activating transcription^3^. However, the NTD’s intrinsic disorder and lack of a well-defined structure present challenges in developing small molecule ligands. Progress has been made in discovering small molecules that interact with the NTD of p53^4,5^ and AR^6,7^, but research on ER-NTD has lagged behind. Recent studies have prominently focused on cyclin-dependent kinase 7 (CDK7)^8-10^, which phosphorylates Ser118 as a prevalent modification in cancerous tumors^11^, enhancing ER activity. Inhibitors of CDK7 are being developed and tested in clinical trials to prevent Ser118 phosphorylation and inhibit ER-NTD activity^10^. Understanding the fundamental disorder-function relationship of ER-NTD is crucial to block its activation effectively.

How a sequence of amino acids determines the structure of disordered proteins is key to decoding their biological function^12-14^. Disordered proteins often undergo post-translational modifications, such as phosphorylation, which can significantly impact their function even with the addition of a single phosphate group. Certain protein kinases, such as MAP kinase or CDK7, can phosphorylate Ser118 within the disordered ER-NTD, activating its transcriptional function^8,15-17^. However, the precise impact of this seemingly simple chemical modification on the conformation and underlying mechanism of functional regulation of ER-NTD has remained elusive. Structural studies of ER-NTD face obstacles due to its highly flexible nature, making conventional tools like crystallography and cryo-EM^18,19^, as well as computational tools like AlphaFold^20,21^, less applicable in capturing its conformational characteristics. Currently, the available biophysical data on ER-NTD is limited^22^, with observations derived from techniques such as circular dichroism^23^, two-dimensional nuclear magnetic resonance (NMR) spectroscopy^23,24^, hydroxyl radical footprinting^25^, and fluorine chemical shift perturbations^26^.

Phosphorylation has been shown to affect long-range interactions in disordered proteins, as observed in the case of p53-NTD^27,28^. Compared to the highly acidic p53-NTD, the ER-NTD has a closer-to-neutral net charge but higher aromatic content, suggesting it may possess more pronounced structural features. However, the lack of sequence-specific resonance assignments has hindered the analysis of NMR data, making it challenging to explore the fundamental sequence-structure-function relationship of this disordered protein.

Here, we identify long-range interactions between distant regions in the ER-NTD amino acid sequence following resonance assignments. We further discover that Ser118 phosphorylation and its phosphomimetic mutation induce conformational changes that extend beyond the immediate vicinity of the phosphorylation site. Unexpectedly, these long-range interactions are not influenced by electrostatic shielding from aqueous salts, despite the negative charges induced by phosphorylation. Instead, our findings demonstrate that hydrophobic interactions, particularly involving amino acids near the phosphorylation site, are crucial in driving these long-range conformational changes. To investigate the underlying mechanism, we introduce single alanine substitutions to modulate the hydrophobic effect of residues such as Phe120 or Leu121, which are adjacent to the phosphorylation site Ser118. Remarkably, these alanine substitutions produce similar long-range conformational influences as those induced by Ser118 phosphorylation and effectively rescue the impaired transcriptional activity observed in a phosphorylation-deficient variant of the NTD and full-length ER. This study establishes a direct connection between the impact of single-site Ser118 phosphorylation on ER-NTD conformations and its functional regulation of transcriptional activation, shedding light on the intricate sequence-structure-function relationship of this disordered protein.

### Conformational contraction and long-range amino acid interactions

We utilized a combination of small-angle X-ray scattering (SAXS) and NMR spectroscopy to investigate the conformational features of the ER-NTD, which consists of a sequence of *N* = 184 residues (Fig. 1A and fig. S1A). To accurately determine its overall dimensions, we conducted size exclusion chromatography-coupled SAXS (SEC-SAXS) measurements (fig. S1B), which improved sample homogeneity compared to our previous flow-cell setup^25^. The observed SEC-SAXS profile (fig. S1C) reveals that the ER-NTD possesses a radius of gyration *R*_*g*_ = 33.9 ± 0.2 Å and a scaling component of *v* = 0.51, as determined through the power-law *R* _*g*_ ∼ *N*^*v*^ analysis^29^ and fitting to an empirically derived molecular form factor^30^. A relavant Kratky plot illustrates the presence of overall structural disorder with modest compaction (Fig. 1B and fig. S1D). Notably, compared to simulated conformations generated from sequence-based flexible-meccano calculations^25^, the pairwise distance distribution derived from SEC-SAXS data (Fig. 1B) appears narrower, suggesting the existence of residual structures and tertiary interactions within the disordered chain.

**Fig. 1.**
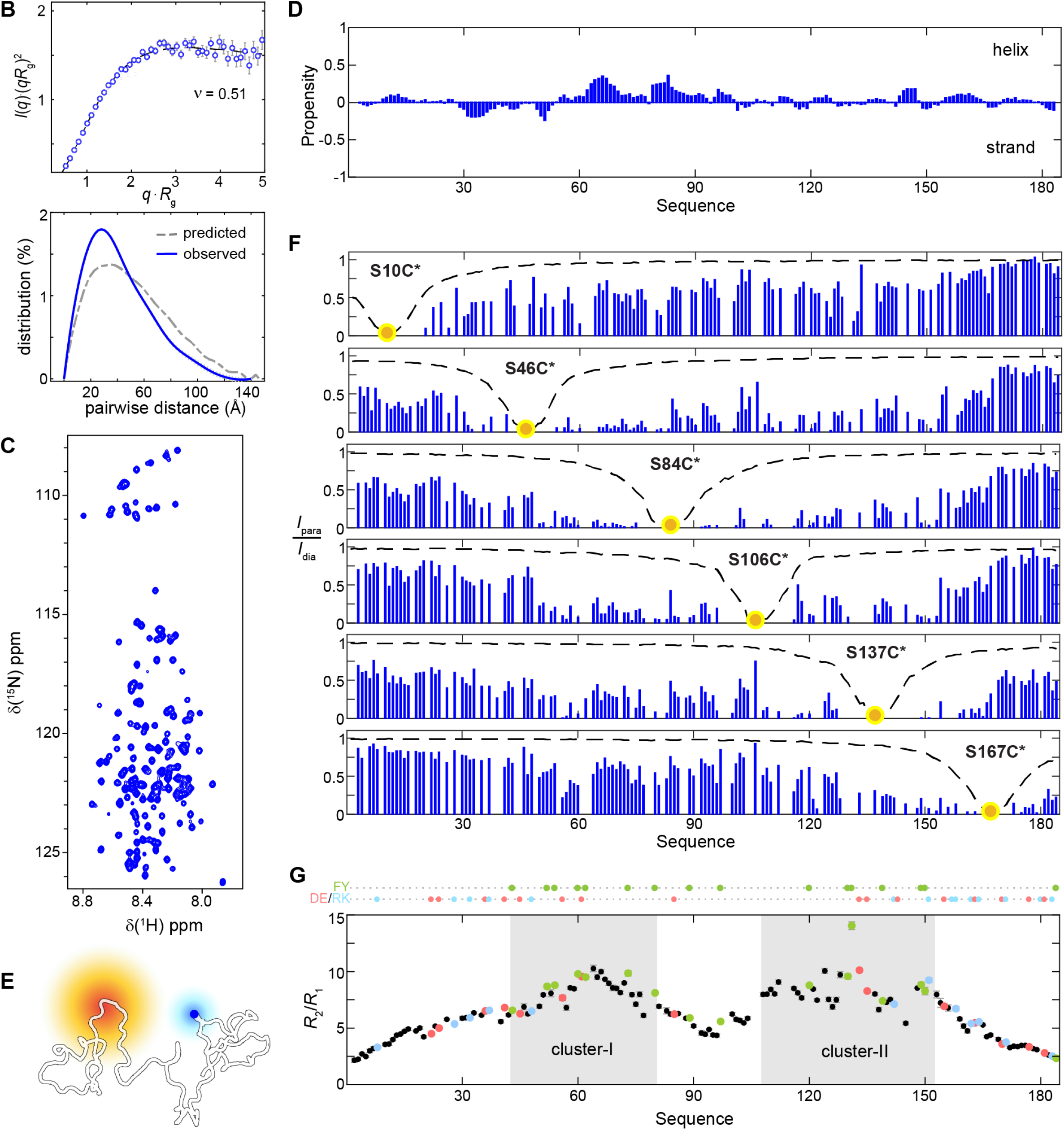
Long-range interactions and hydrophobic clustering of residual structures. (**A**) ER-NTD sequence decorated with negatively charged (light red), positively charged (light blue), and aromatic residues (light green). Ser118, highlighted with a red box. (**B**) Kratky plot for SEC-SAXS data of ER-NTD collected at 4°C using a pre-cooled running buffer buffer for size-exclusion chromatography and related pairwise distance distribution. Dashed line, calculated distribution from conformers generated via flexible-meccano simulations^31^. (**C**) ^1^H-^15^N HSQC spectrum. Resonance assignments are shown in fig. S2. (**D**) Secondary structure propensity (SSP), calculated using ^1^H, ^15^N, ^13^C (C^α^, C^β^, and C^′^) chemical shifts^32^. Low SSP scores indicate an overall lack of persistent secondary structure. (**E**) Schematic of the paramagnetic effect between a nitroxide spin (yellow) and an observed NMR-active nucleus (blue). (**F**) PRE profiles of amide protons after MTSL spin labeling of six cysteine mutants, S10C, S46C, S84C, S106C, S137C, and S167C, each analyzed by ^1^H-^15^N HSQC spectra for a protein concentration of 30 μM. The intensity ratio between the paramagnetic form (*I*_para_) and the diamagnetic form (*I*_dia_), representing the presence and absence of the active spin label, provides distance information between amide protons and the spin label, with the ratio values ranging from 0 (close) to 1 (far). Yellow circle, the position of spin labeling; dashed line, calculated from flexible-meccano simulations^31^. (**G**) ^15^N amide relaxation rates using *R*_2_/*R*_1_ ratios reveal two clusters of residues, 43 to 80 (cluster-I) and 108 to 151 (cluster-II). Color dot, charged (positive in blue; negative in red), and aromatic (green) amino acid. HSQC spectra were recorded using 30 μM protein samples at pH 7.4 in a buffer containing 20 mM sodium phosphate, 100 mM NaCl, 0.5 mM EDTA, 0.1 mM PMSF, and 5% (v/v) D_2_O, at 850 MHz and 4°C unless stated otherwise.

We next conducted NMR studies to investigate the contribution of the ER-NTD amino acids to the observed conformational contraction. The ^1^H-^15^N heteronuclear single-quantum coherence (HSQC) spectrum showed a narrow proton chemical shift dispersion, characteristic of intrinsically disordered proteins (Fig. 1C). Resonance assignments (fig. S2) and resulting C_α_ and C_β_ chemical shifts indicated that the ER-NTD has minimal persistent secondary structure, except for two short alanine-rich segments (i.e., A^64^AAAA^68^ and A^86^AA^88^) exhibiting transient helical features with an approximate propensity of 30% (Fig. 1D), resembling those observed in AR-NTD^7^. Compared to previously reported 2D-NMR spectra without resonance assignments^23,24^, including our initial data^25^, the newly obtained NMR spectra exhibit well-resolved cross-peaks, providing a residue-specific level of detail in characterizing the ER-NTD conformational features.

To probe long-range interactions in ER-NTD, we conducted paramagnetic relaxation enhancement (PRE) measurements, which are sensitive to distances between a nitroxide spin label and nearby amino acids within a range of approximately 25 Å^33,34^ (Fig. 1E). As an advantage, ER-NTD lacks native cysteine residues (Fig. 1A), so we introduced a cysteine residue at each comparable serine site, allowing its attachment with an MTSL spin-label. The intensity ratios between the paramagnetic and diamagnetic forms converged when comparing two doubling protein concentrations (fig. S3), indicating that the observed ratios predominantly originated from intramolecular interactions, with minimal intermolecular interactions.

The paramagnetic effects demonstrated variability across the six labeling positions (Fig. 1F). Analysis of the PRE profiles revealed low-intensity ratios for residues 40 to 150 within mid-sequence regions. This attenuation in intensity demonstrates significant long-range interactions in this region. These findings align with previous observations in the disordered p53-NTD^27,28^ and other chemically denatured or unfolded proteins^35-38^. The reduction in the paramagnetic effect was consistently broad and supported by ^15^N relaxation rate measurements (longitudinal *R*_1_ and transverse *R*_2_). Despite the absence of persistent secondary structure, two distinct regions with larger *R*_2_/*R*_1_ ratios were identified (Fig. 1G and fig. S4A), indicating enhanced relaxation and restricted motion^39,40^. These regions, spanning approximately residues 43 to 80 (cluster-I) and residues 108 to 151 (cluster-II), exhibited slower internal motion compared to the apparent random chain rotational correlation time (fig. S4B). These residual structures likely arise from electrostatic interactions between charged amino acids within the two regions or the transient clustering of hydrophobic amino acids, particularly considering the high content of aromatic residues.

### Phosphorylation-induced long-range conformational changes

To investigate the impact of Ser118 phosphorylation on the disordered conformation of ER-NTD, we used *in vitro* phosphorylated ER-NTD by MAP kinase^41^ (MAPK1, also known as ERK2; fig. S5) and a phosphomimetic S118D mutant. Interactions with peptidylprolyl isomerase Pin1 (fig. S6), which is known for its exclusive binding to the phosphorylated Ser118/Pro119 motif^24^, provided evidence for the use of Asp at position 118, *i*.*e*., S118D, as a phosphomimetic mutation. Previous studies have shown that Ser118 phosphorylation by MAP kinase or CDK7 enhances ER-mediated reporter activity^8,15^. Consistent with these findings, we observed that the phosphomimetic S118D variant exhibited increased ER activity, showing 2-3 fold higher activity than the wild-type protein in HEK293 and MCF7 cells (fig. S7A and fig. S7B). Importantly, we also observed enhanced activity by S118D in full-length ER (fig. S7C). In contrast, the phosphorylation-deficient S118A variant exhibited only 50-60% of the wild-type activity. These results indicate that Ser118 phosphorylation and its phosphomimetic S118D mutant contribute to the transcriptional activity of both the NTD and full-length ER. These observations suggest that Ser118 phosphorylation modulates the disordered conformation of ER-NTD.

To determine the specific impact of phosphorylation on the disordered conformation, we conducted resonance assignments for Ser118 phosphorylation (pS118) and the phosphomimetic S118D variant. We observed that both pS118 and S118D trigger long-range changes that extend beyond its immediate local surroundings (Fig. 2A and fig. S8), affecting amino acid sites Glu56 and Gly57 that are far apart in sequence (Fig. 2B and fig. S9). However, we found little or no change in secondary structure propensity (Fig. 2C and fig. S10), hydrophobic clustering of residual structures (Fig. 2C and fig. S11). Additionally, we investigated the impact of Ser118 phosphorylation on solvent accessibility by probing amide proton relaxation rates Δ*R*_2_(^1^H^N^) as a function of an increasing concentration series of soluble paramagnetic agents gadodiamide^42-44^ and found no significant alteration in the residue-specific solvent accessibility upon Ser118 phosphorylation (fig. S12). Since the positions of pSer118 and Glu56/Gly57 are not close together in sequence, we can infer that these phosphorylation-induced changes in the chemical shift of Glu56 and Gly57 are primarily due to long-range tertiary modulation between clusters of amino acids, rather than local changes within the clusters.

**Fig. 2.**
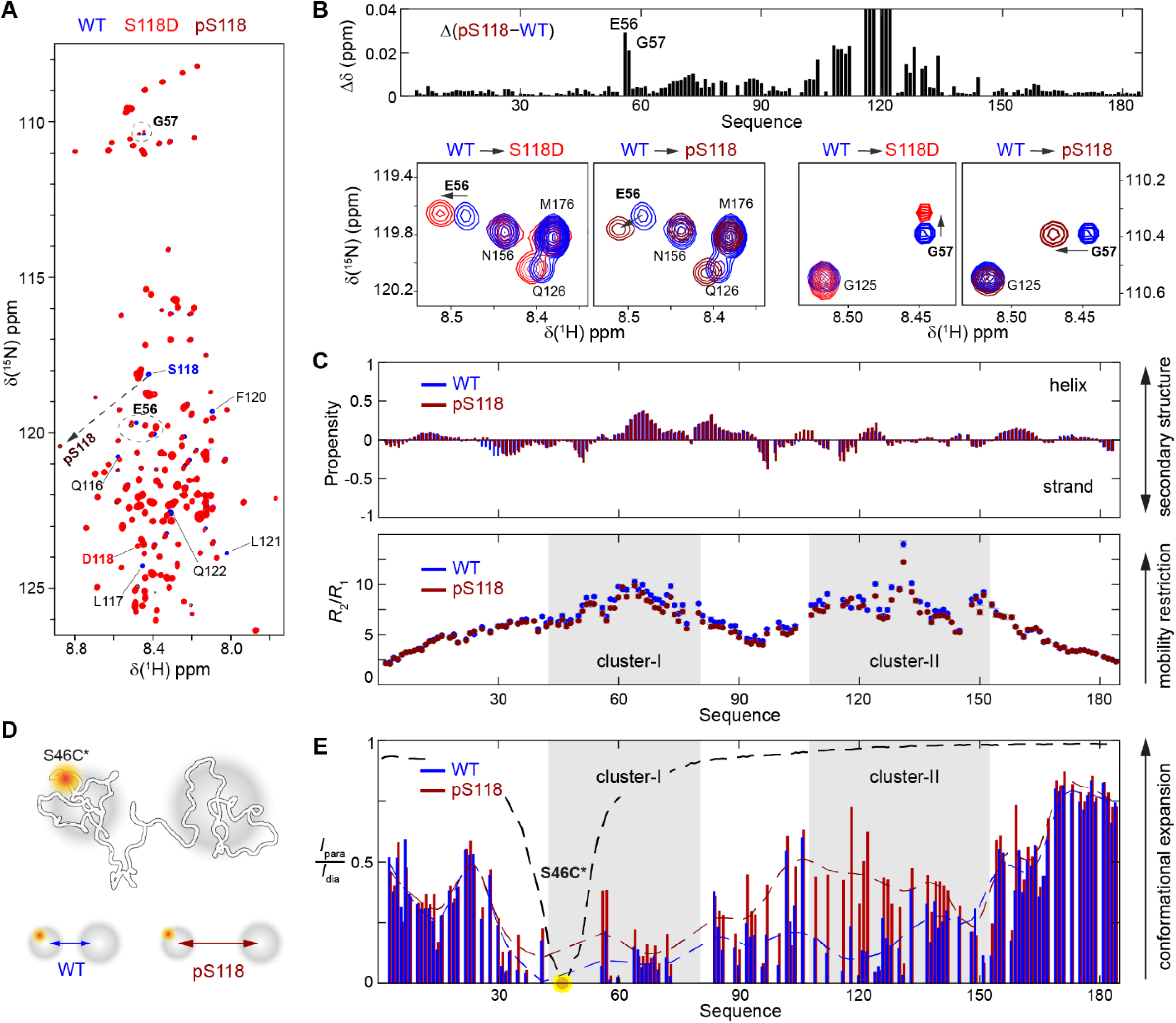
Phosphorylation of Ser118 triggers long-range conformational changes. (**A**) Comparison of ^1^H-^15^N HSQC spectra reveals the difference between wild-type (blue) and the S118D (red) and pSer118 (dark red) variants. Annotated residues display notable differences and circled are magnified in (B). (**B)** Chemical shift changes of backbone ^5^N and ^1^H upon Ser118 phosphorylation (also fig. S9). Bottom, residues Glu56 (left) and Gly57 (right) with notable changes in chemical shift induced by S118D and pSer118, respectively. (**C**) Comparison of secondary structure propensity calculated using ^1^H, ^5^N, ^13^C (C^α^, C^β^, and C^’^) chemical shifts (top) and ^5^N amide relaxation rate *R*_2_/*R*_1_ ratios (bottom) between the wild-type and the pSer118 variant. (**D** and **E**) Ser118 phosphorylation increases the physical distancing between two hydrophobic clusters, as observed using S46C spin-labeling (D) and demonstrated in the PRE profiles of both the wild-type and pSer118 variant (E). Yellow circle, the position of spin labeling; dash lines, visual aid. All HSQC spectra were recorded at 850 MHz and 4°C.

We subsequently used PRE measurements to investigate the impact of Ser118 phosphorylation on long-range interactions between the two clusters of residual structures as probed by ^5^N amide relaxation measurements (Fig. 1G). To minimize any potential perturbations introduced by spin labeling that may affect the clustering, we deliberately selected the S46C site located at the edge of cluster-I, which is also away from cluster-II and Ser118 (Fig. 2D). Strikingly, we observed a systematic increase in the PRE profile of cluster-II, particularly in the vicinity of Ser118 (Fig. 2E). This observation indicates the significant impact of Ser118 phosphorylation on long-range interactions between the two clusters. Specifically, phosphorylation of Ser118 results in the two clusters moving away, expanding the disordered conformation of ER-NTD (Fig. 2D). This increase in physical distancing is further supported by another spin labeling at S84C in both the S118D and pS118 variants (fig. S13), albeit to a lesser extent due to its closer proximity to both clusters when compared to S46C spin labeling (Fig. 2E). These paramagnetic effects on a group of amino acids collectively align with the overall expansion observed in the global conformation, as revealed by SEC-SAXS data, particularly with a 3-Å increase in Rg (fig. S14) observed upon the phosphomimetic S118D mutation. These findings demonstrate that Ser118 phosphorylation increases the separation between two clusters of hydrophobic amino acids and shifts the conformational ensemble toward more open conformations while preserving the clusters of local residual structures.

### Hydrophobic interactions drive phosphorylation-induced conformational changes

We conducted further investigations to elucidate the mechanistic basis of phosphorylation-induced conformational changes in the disordered ER-NTD. Phosphorylation of Ser118 has two specific effects stemming from its phosphoryl group (Fig. 3A): the introduction of negative charges that may lead to electrostatic repulsion with other negatively charged residues within the clusters (Fig. 2C) and the alteration of local hydrophobicity resulting in a less hydrophobic environment and weakening long-range interactions with other hydrophobic residues within the clusters. To specifically test the possibility of electrostatic repulsion, we increased the salt concentration to determine if these interactions could be shielded. Our analyses of chemical shift perturbations revealed minimal conformational changes in the region containing Glu56 and Gly57 when the salt concentration was varied. This observation remained consistent except for the termini regions, which are rich in charged residues such as those near Lys32 and Arg158 (Fig. 3B and Fig. 3C). Importantly, these termini regions are located outside the hydrophobic clusters, and the observed chemical shift perturbations in these regions primarily arise from electrostatic interactions that are shielded by the presence of salt (Fig. 3C). Thus, the most significant conformational changes observed in response to Ser118 phosphorylation are not driven by electrostatic interactions from the added negative charges of the phosphoryl group. Instead, these changes can be attributed to long-range hydrophobic interactions involving amino acids near or including the phosphorylation site.

**Fig. 3.**
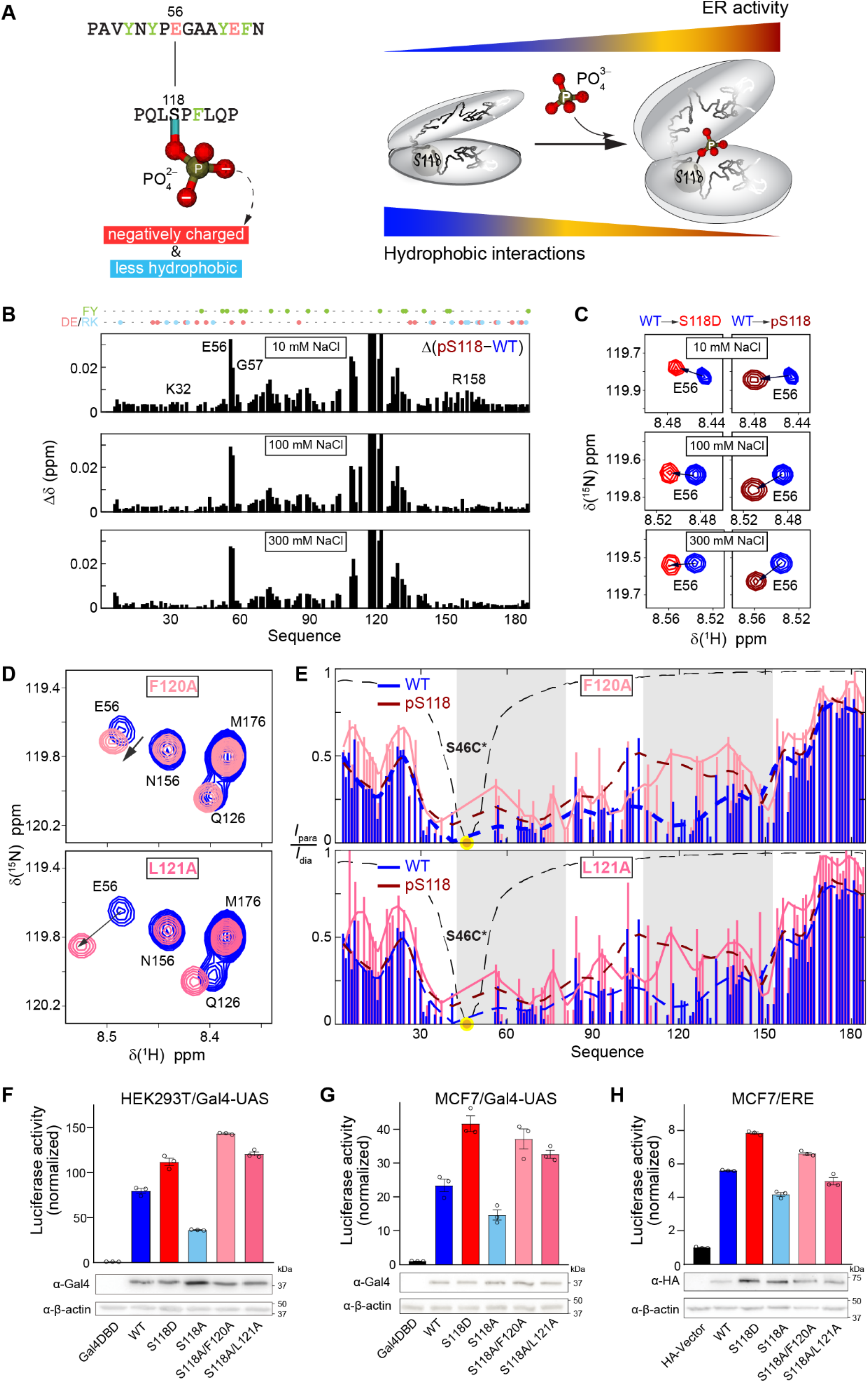
Hydrophobic interactions drive phosphorylation-induced conformational changes and ER transcriptional activity. (**A**) Long-range interactions between the segments involving pSer118 and Glu56 (left), as depicted by the structure-function regulation of Ser118 phosphorylation on ER activity (right). Ball-and-stick, phosphoryl group of pS118. Aromatic residue, light green; charged residues, light red. (**B**) Chemical shift changes Δδ for ^1^H and ^5^N induced by Ser118 phosphorylation measured at different NaCl concentrations (10, 100, and 300 mM). (**C**) Magnified regions of ^1^H-^5^N HSQC spectra displaying the chemical shift change of Glu56 for the S118D (left) and pSer118 (right) variants, as observed under three different salt conditions relative to the wild-type. (**D**) Chemical shift changes of Glu56 upon the F120A or L121A mutation, compared to the wild-type (blue). All HSQC spectra were recorded at 850 MHz and 4°C. (**E**) F120A or L121A induces long-range conformational changes, resembling the effect of Ser118 phosphorylation (dark red line), as assessed through PRE measurements using S46C spin labeling, compared to the wild-type (blue). Yellow circle, the position of spin labeling; dash/solid lines, visual aid. (**F** and **G**) F120A or L121A rescues the transcription activity of ER-NTD impaired by S118A in HEK293T cells (F) and MCF7 cells (G), expressing WT, S118D, and S118A proteins. (**H**) F120A or L121A rescues the transcription activity of the full-length ER that was impaired by S118A, using MCF7 cells that have been treated with 100 nM17β-estradiol for 24 hours. Transient transfections and dual-luciferase reporter assays were conducted. Mean ± SEM, calculated from three biological repeats. Bottom, western blot analysis of protein levels.

To further investigate the hypothesis that Ser118 phosphorylation induces long-range changes through hydrophobic interactions, we introduced alanine substitutions at positions Phe120 or Leu121. These positions were chosen due to their proximity to Ser118. By introducing these alanine mutations, we aimed to mimic the reduction in hydrophobicity that occurs upon phosphorylation. The chemical shifts of the Glu56/Gly57 region were used as an indicator to monitor the effect of these alanine substitutions (Fig. 2B). Our results demonstrate that both the F120A and L121A mutations induce chemical shift changes (Fig. 3D), similar to those observed in S118D and pSer118 variants (Fig. 2B). Remarkably, both F120A and L121A affect paramagnetic intensity attenuation and increase the physical distancing between the two hydrophobic clusters, as assessed by the S46 spin labeling technique (Fig. 3E), closely resembling the conformational alternation of Ser118 phosphorylation (Fig. 3E). These findings provide direct validation of the hydrophobic-driven hypothesis and establish the involvement of hydrophobic interactions in the conformational changes induced by Ser118 phosphorylation.

These results also challenge the commonly assumed mechanism of phosphorylation-induced conformational changes, which predominantly rely on electrostatic interactions between charged amino acids.

### Reduction of hydrophobic interactions rescues ER transcriptional activity

Given the wild-type’s susceptibility to phosphorylation in cellular contexts, we selected the phosphorylation-deficient S118A variant as a reference to investigate the impact on the activation function. To mimic the reduction in hydrophobicity caused by phosphorylation, we introduced alanine substitutions at positions Phe120 or Leu121, generating two double-mutants, S118A/F120A and S118A/L121A. These mutations were then incorporated into luciferase reporter assays to assess their functional effects. Previous studies have established that phosphorylation of Ser118 by MAP kinase or CDK7 enhances the transcriptional activity of the NTD and full-length ER proteins^8,15^. Consistent with these findings, we observed a similar increase in reporter activity with the phosphomimetic S118D variant and a reduction with the phosphorylation-deficient S118A mutant (fig. S7). Strikingly, in HEK293 and MCF7 cells, both F120A and L121A rescued the largely abolished transcriptional activity caused by S118A (Fig. 3F and Fig. 3G). Furthermore, in the context of full-length ER, the two double mutants S118A/F120A and S118A/L121A exhibited activity comparable to the phosphomimetic S118D variant (Fig. 3H). These findings underscore the functional importance of hydrophobic-driven modulation in ER activity, emphasizing the significance of hydrophobic interactions, *independent of phosphorylation*, in the activation function of ER for its transcriptional activity.

The functional rescue observed, along with the corresponding conformational changes (Fig. 3D and Fig. 3E), provides strong evidence that precise modulation of long-range interactions in the vicinity of Ser118 is crucial for releasing the disordered ER-NTD from its inactive wild-type state and promoting ER activity through Ser118 phosphorylation. Owing to the weak and long-range nature of these residual structures, even minor changes in the local environment, such as Ser118 phosphorylation or the introduction of F120A/L121A mutations, have a significant impact on shifting the conformational equilibrium towards active ER states (Fig. 3A). These observations shed light on the mechanisms underlying ER disorder-activity interplay in a disordered protein, emphasizing the crucial role of long-range hydrophobic interactions in regulating the transcriptional function of this disordered ER-NTD.

## Discussion

Post-translational modifications, particularly phosphorylation, are pivotal for the functional regulation of disordered proteins^12,45^. However, it has been challenging to connect specific phosphorylation events with the function regulation of disordered proteins, such as the phosphorylation of Ser118 (pS118) within its disordered ER-NTD. ER-pS118 is frequently observed in cancerous tumors^11^ and associated with transcriptional activation^8,15^, but the precise mechanism linking this single-site phosphorylation to the regulation of its activation function has remained unclear. Our investigation reveals that Ser118 phosphorylation expands the conformational ensemble of ER-NTD by reducing long-range interactions between distant regions in the amino acid sequence, shifting it towards more open conformational states during activation (Fig. 3A). These conformational changes occur through long-range interactions between the distant regions in the amino acid sequence primarily facilitated by the hydrophobic clustering of residual structures. These findings establish a new sequence-structure-function paradigm in disordered proteins, demonstrating how single-site phosphorylation, specifically Ser118 phosphorylation, exerts its influence on ER activity.

Long-range hydrophobic interactions between clusters of aromatic residues were observed in residual structures of chemically denatured or unfolded lysozyme, and these clusters are disrupted when the bulky hydrophobic tryptophan residue at position 62 is substituted with glycine (W62G)^39^. In the case of ER-NTD, phosphorylation of Ser118 leads to an increased physical distancing between hydrophobic clusters of residual structures while maintaining residual hydrophobic clustering. Compared to the W62G mutation of lysozyme, Ser118 phosphorylation in ER-NTD results in a similar decrease in hydrophobicity^46,47^. However, phosphorylation of Ser118 reduces hydrophobicity and leads to the expansion of ER-NTD conformation while activating ER transcriptional function. These findings also align with studies on p53-NTD, where phosphorylation caused the dissociation of two amino acid segments, a phenomenon that cannot be fully explained by electrostatic repulsion resulting from the added negative charges of the phosphoryl group^28^. Remarkably, even without invoking phosphorylation itself, alanine substitution of nearby hydrophobic amino acids can produce conformational alterations similar to those induced by phosphorylation and restore the impaired activation function caused by a phosphorylation-deficient mutation. This study thus fills a knowledge gap by elucidating how single-site phosphorylation modulates the conformational ensemble and influences the activation function through the newfound sequence-structure-function paradigm of this disordered protein.

Considering the inherent flexibility of disordered proteins like ER-NTD, which engage in weak and promiscuous interactions with a wide range of partner proteins^12,48^, the fine-tuning mediated by Ser118 phosphorylation has the potential to tilt the delicate balance within the conformational ensemble of ER-NTD and beyond. This phosphorylation event can propagate its impact, regulating downstream protein-protein interactions. The downstream regulation of such an extensive protein-protein interacting network is important for understanding the functional implications of small-molecule inhibitors that can effectively block the transcriptional activation of disordered ER-NTD as a target of interest in drug discovery.

## Notes

### Competing Interest Statement

The authors have declared no competing interest.

## Reference

1. Lambert, S.A., et al. The Human Transcription Factors. Cell 175, 598–599 (2018).

2. Darnell, J.E., Jr. Transcription factors as targets for cancer therapy. Nat Rev Cancer 2, 740–749 (2002).

3. Henley, M.J. & Koehler, A.N. Advances in targeting ‘undruggable’ transcription factors with small molecules. Nat Rev Drug Discov 20, 669–688 (2021).

4. Ruan, H., et al. Computational strategy for intrinsically disordered protein ligand design leads to the discovery of p53 transactivation domain I binding compounds that activate the p53 pathway. Chem Sci 12, 3004–3016 (2020).

5. Zhao, J., et al. EGCG binds intrinsically disordered N-terminal domain of p53 and disrupts p53-MDM2 interaction. Nat Commun 12, 986 (2021).

6. Andersen, R.J., et al. Regression of castrate-recurrent prostate cancer by a small-molecule inhibitor of the amino-terminus domain of the androgen receptor. Cancer cell 17, 535–546 (2010).

7. De Mol, E., et al. EPI-001, A Compound Active against Castration-Resistant Prostate Cancer, Targets Transactivation Unit 5 of the Androgen Receptor. ACS Chem Biol 11, 2499–2505 (2016).

8. Chen, D., et al. Activation of estrogen receptor alpha by S118 phosphorylation involves a ligand-dependent interaction with TFIIH and participation of CDK7. Molecular cell 6, 127–137 (2000).

9. Patel, H., et al. ICEC0942, an Orally Bioavailable Selective Inhibitor of CDK7 for Cancer Treatment. Mol Cancer Ther 17, 1156–1166 (2018).

10. Sava, G.P., Fan, H., Coombes, R.C., Buluwela, L. & Ali, S. CDK7 inhibitors as anticancer drugs. Cancer Metastasis Rev 39, 805–823 (2020).

11. Cancer Genome Atlas Research, N., et al. The Cancer Genome Atlas Pan-Cancer analysis project. Nat Genet 45, 1113–1120 (2013).

12. Wright, P.E. & Dyson, H.J. Intrinsically disordered proteins in cellular signalling and regulation. Nature reviews. Molecular cell biology 16, 18–29 (2015).

13. Marsh, J.A. & Forman-Kay, J.D. Sequence determinants of compaction in intrinsically disordered proteins. Biophys J 98, 2383–2390 (2010).

14. Uversky, V.N., Gillespie, J.R. & Fink, A.L. Why are “natively unfolded” proteins unstructured under physiologic conditions? Proteins 41, 415–427 (2000).

15. Kato, S., et al. Activation of the estrogen receptor through phosphorylation by mitogen-activated protein kinase. Science 270, 1491–1494 (1995).

16. Le Goff, P., Montano, M.M., Schodin, D.J. & Katzenellenbogen, B.S. Phosphorylation of the human estrogen receptor. Identification of hormone-regulated sites and examination of their influence on transcriptional activity. J Biol Chem 269, 4458–4466 (1994).

17. Ali, S., Metzger, D., Bornert, J.M. & Chambon, P. Modulation of transcriptional activation by ligand-dependent phosphorylation of the human oestrogen receptor A/B region. Embo J 12, 1153–1160 (1993).

18. Yi, P., et al. Structure of a biologically active estrogen receptor-coactivator complex on DNA. Molecular cell 57, 1047–1058 (2015).

19. Yi, P., et al. Structural and Functional Impacts of ER Coactivator Sequential Recruitment. Molecular cell 67, 733–743 e734 (2017).

20. Jumper, J., et al. Highly accurate protein structure prediction with AlphaFold. Nature 596, 583–589 (2021).

21. Ruff, K.M. & Pappu, R.V. AlphaFold and Implications for Intrinsically Disordered Proteins. J Mol Biol 433, 167208 (2021).

22. Luo, S., Wohl, S., Zheng, W. & Yang, S. Biophysical and Integrative Characterization of Protein Intrinsic Disorder as a Prime Target for Drug Discovery. Biomolecules 13, 530 (2023).

23. Warnmark, A., Wikstrom, A., Wright, A.P.H., Gustafsson, J.A. & Hard, T. The N-terminal regions of estrogen receptor alpha and beta are unstructured in vitro and show different TBP binding properties. J Biol Chem 276, 45939–45944 (2001).

24. Rajbhandari, P., et al. Regulation of estrogen receptor alpha N-terminus conformation and function by peptidyl prolyl isomerase Pin1. Mol Cell Biol 32, 445–457 (2012).

25. Peng, Y., et al. A Metastable Contact and Structural Disorder in the Estrogen Receptor Transactivation Domain. Structure 27, 229–240 e224 (2019).

26. Zheng, W., Du, Z., Ko, S.B., Wickramasinghe, N.P. & Yang, S. Incorporation of D(2)O-Induced Fluorine Chemical Shift Perturbations into Ensemble-Structure Characterization of the ERalpha Disordered Region. J Phys Chem B 126, 9176–9186 (2022).

27. Vise, P., Baral, B., Stancik, A., Lowry, D.F. & Daughdrill, G.W. Identifying long-range structure in the intrinsically unstructured transactivation domain of p53. Proteins 67, 526–530 (2007).

28. Lum, J.K., Neuweiler, H. & Fersht, A.R. Long-range modulation of chain motions within the intrinsically disordered transactivation domain of tumor suppressor p53. J Am Chem Soc 134, 1617–1622 (2012).

29. Flory, P.J. Principles of Polymer Chemistry, (Cornell University Press, Ithaca and London, 1953).

30. Riback, J.A., et al. Innovative scattering analysis shows that hydrophobic disordered proteins are expanded in water. Science 358, 238–241 (2017).

31. Ozenne, V., et al. Flexible-meccano: a tool for the generation of explicit ensemble descriptions of intrinsically disordered proteins and their associated experimental observables. Bioinformatics 28, 1463–1470 (2012).

32. Tamiola, K., Acar, B. & Mulder, F.A. Sequence-specific random coil chemical shifts of intrinsically disordered proteins. J Am Chem Soc 132, 18000–18003 (2010).

33. Battiste, J.L. & Wagner, G. Utilization of site-directed spin labeling and high-resolution heteronuclear nuclear magnetic resonance for global fold determination of large proteins with limited nuclear overhauser effect data. Biochemistry 39, 5355–5365 (2000).

34. Tang, C., Iwahara, J. & Clore, G.M. Visualization of transient encounter complexes in protein-protein association. Nature 444, 383–386 (2006).

35. Cordeiro, T.N., et al. Interplay of Protein Disorder in Retinoic Acid Receptor Heterodimer and Its Corepressor Regulates Gene Expression. Structure 27, 1270–1285 e1276 (2019).

36. Lietzow, M.A., Jamin, M., Dyson, H.J. & Wright, P.E. Mapping long-range contacts in a highly unfolded protein. J Mol Biol 322, 655–662 (2002).

37. Salmon, L., et al. NMR characterization of long-range order in intrinsically disordered proteins. J Am Chem Soc 132, 8407–8418 (2010).

38. Bertoncini, C.W., et al. Release of long-range tertiary interactions potentiates aggregation of natively unstructured alpha-synuclein. Proc Natl Acad Sci U S A 102, 1430–1435 (2005).

39. Klein-Seetharaman, J., et al. Long-range interactions within a nonnative protein. Science 295, 1719–1722 (2002).

40. Martin, E.W., et al. Valence and patterning of aromatic residues determine the phase behavior of prion-like domains. Science 367, 694–699 (2020).

41. Likhite, V.S., Stossi, F., Kim, K., Katzenellenbogen, B.S. & Katzenellenbogen, J.A. Kinase-specific phosphorylation of the estrogen receptor changes receptor interactions with ligand, deoxyribonucleic acid, and coregulators associated with alterations in estrogen and tamoxifen activity. Mol Endocrinol 20, 3120–3132 (2006).

42. Gong, Z., Schwieters, C.D. & Tang, C. Theory and practice of using solvent paramagnetic relaxation enhancement to characterize protein conformational dynamics. Methods 148, 48–56 (2018).

43. Hocking, H.G., Zangger, K. & Madl, T. Studying the structure and dynamics of biomolecules by using soluble paramagnetic probes. Chemphyschem : a European journal of chemical physics and physical chemistry 14, 3082–3094 (2013).

44. Kooshapur, H., Schwieters, C.D. & Tjandra, N. Conformational Ensemble of Disordered Proteins Probed by Solvent Paramagnetic Relaxation Enhancement (sPRE). Angew Chem Int Ed Engl 57, 13519–13522 (2018).

45. Bah, A., et al. Folding of an intrinsically disordered protein by phosphorylation as a regulatory switch. Nature 519, 106–U240 (2015).

46. Kapcha, L.H. & Rossky, P.J. A simple atomic-level hydrophobicity scale reveals protein interfacial structure. J Mol Biol 426, 484–498 (2014).

47. Perdikari, T.M., et al. A predictive coarse-grained model for position-specific effects of post-translational modifications. Biophys J 120, 1187–1197 (2021).

48. Kim, M., et al. A protein interaction landscape of breast cancer. Science 374, eabf3066 (2021).

