## Supplemental Figures for "Phosphorylation modulates estrogen receptor disorder by altering long-range hydrophobic interactions"

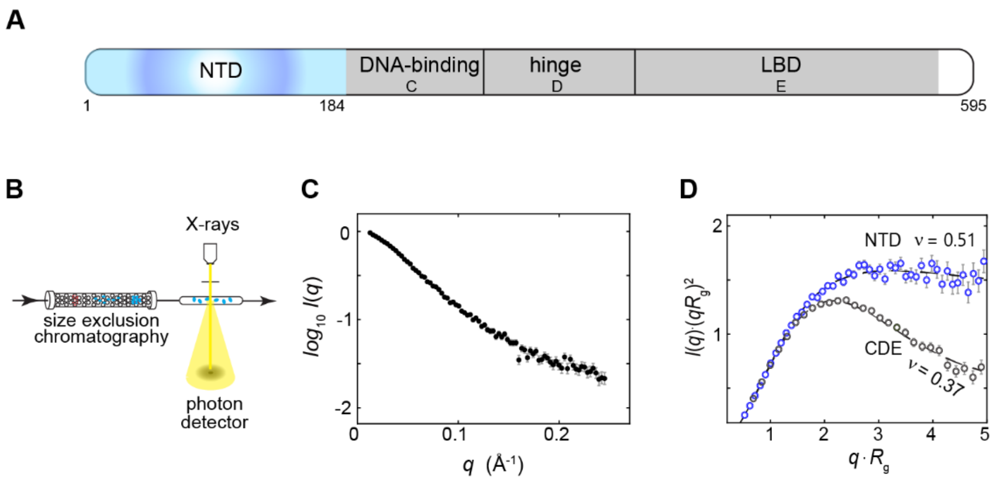


**Fig. S1. The structural disorder observed in ER-NTD via SEC-SAXS.**

(**A**) Modular domains of the ER, encompassing the N-terminal domain (NTD), DNA-binding domain (DBD) or C domain, domain-connecting hinge region or D domain, and a C-terminal ligand-binding domain (LBD) or E domain.

(**B**) Schematic representation of the SEC-SAXS data acquisition process.

(**C**) SEC-SAXS data of ER-NTD acquired at 4°C using a pre-cooled running buffer for size-exclusion chromatography. $I(q)$, scattering intensity with error bars (normalized by zero-angle scattering); $q$, the amplitude of the X-ray scattering vector relating to the scattering angle.

(**D**) Kratky plot for the SEC-SAXS data of ER-NTD (in blue), compared to the folded DBD-hinge-LBD (marked as CDE in gray). The SEC-SAXS data of ER-CDE were taken from Ref.[1]. $v$, a scaling component of the $R_g \sim N^v$ power-law analysis, with $R_g$ denoting the radius of gyration and $N$ representing the number of residues.

The SEC-SAXS data of wild-type ER-NTD are available in the Small Angle Scattering Biological Databank.

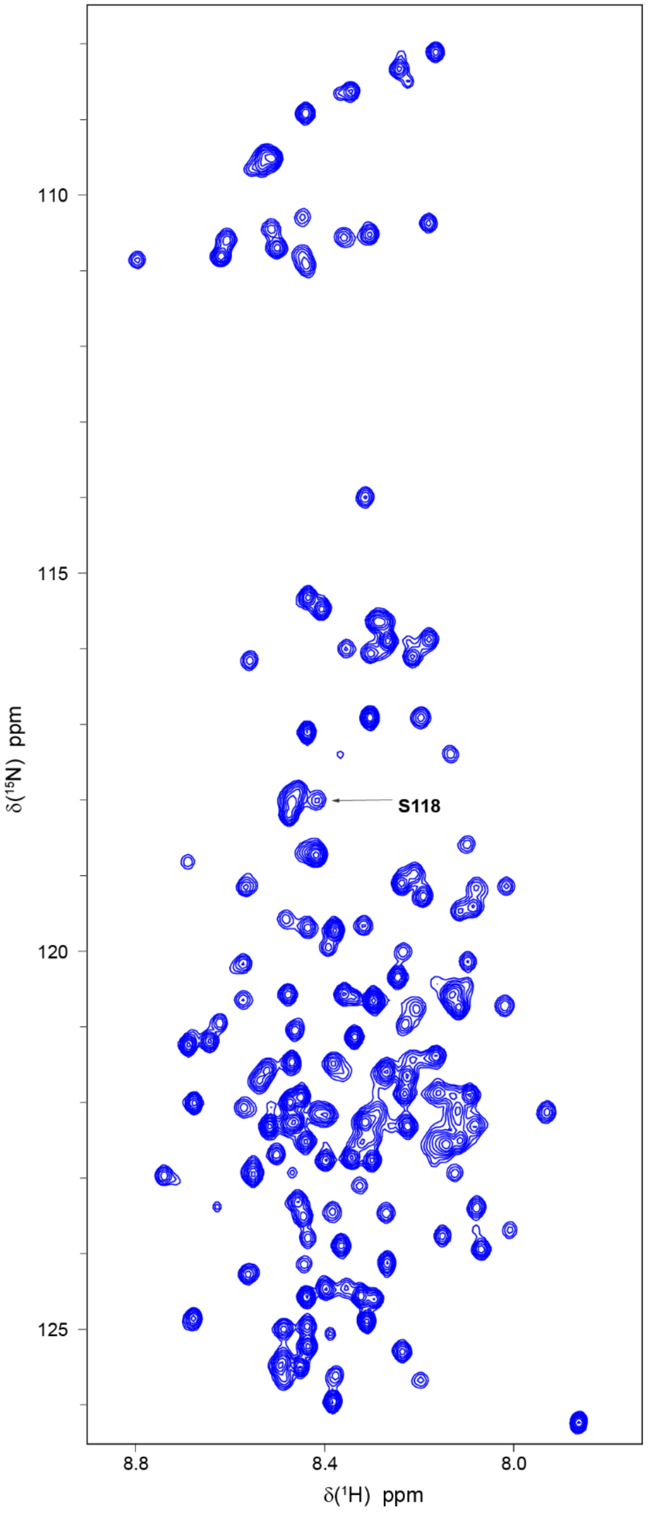


**Fig. S2. Resonance assignments of wild-type ER-NTD depicted on the $^1H$-$^{15}N$ HSQC spectrum**. Deposited assignments of wild-type ER-NTD are available in the Biomolecular NMR database.

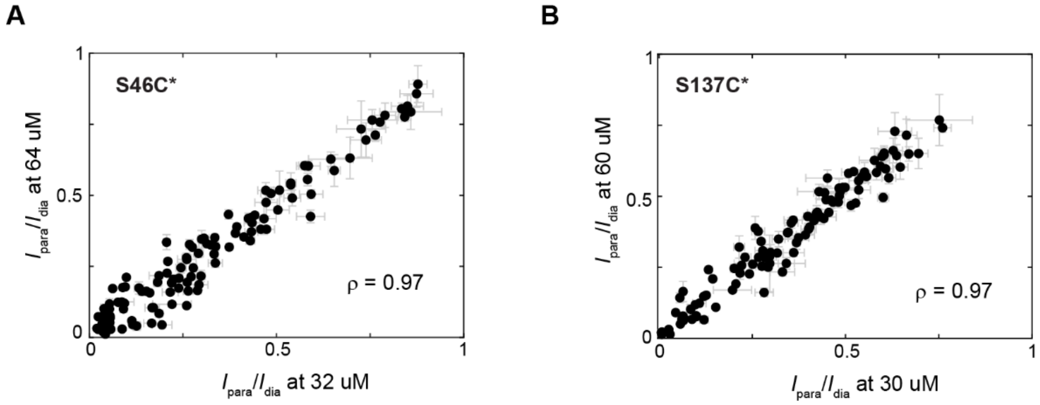


**Fig. S3. Correlation of PRE intensity ratios at two doubling ER-NTD concentrations.**

The intensity ratios between the paramagnetic form ($I_{\text{para}}$) and the diamagnetic ($I_{\text{dia}}$) form were examined and compared between 32 μM and 64 μM for the S46C spin labeling (**A**) and between 30 μM and 60 μM for the S137C spin labeling (**B**). The intensity ratios converge between the two doubling protein concentrations, as determined by their Pearson correlation coefficient (ρ) values.

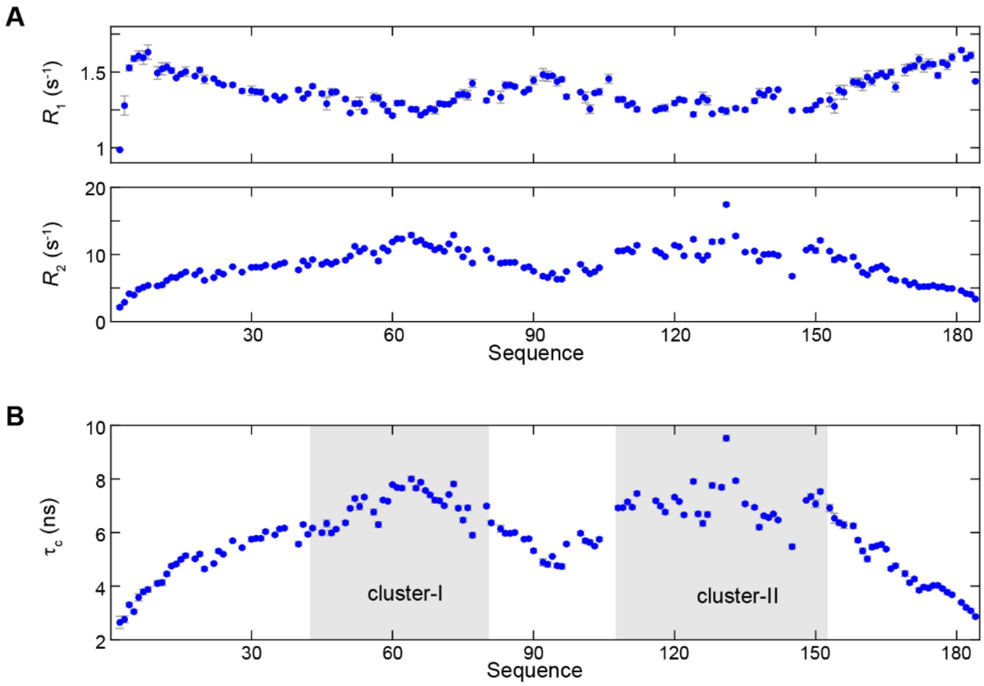


**Fig. S4. $^{15}$N relaxation measurements of wild-type ER-NTD.**

**(A)** The $^{15}$N-$R_1$ and $^{15}$N-$R_2$ profiles at 850 MHz.

**(B)** Overall correlation time ($\tau_c$) along the amino acid sequence, as determined from the measured $^{15}$N relaxation rates.

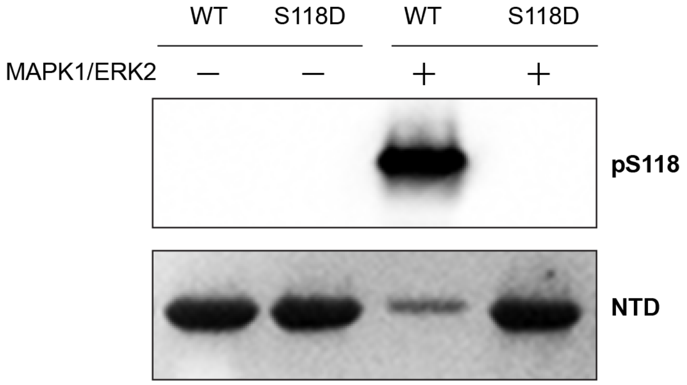


**Fig. S5.** ***In vitro*** **phosphorylation of Ser118 by MAP kinase MAPK1/ERK2.**

Western blot analysis for the phosphorylation status of Ser118 (pS118) using an anti-phospho-ER-Ser118 antibody, with S118D as a negative control. The total NTD proteins were visualized through staining with Ponceau S staining solution.

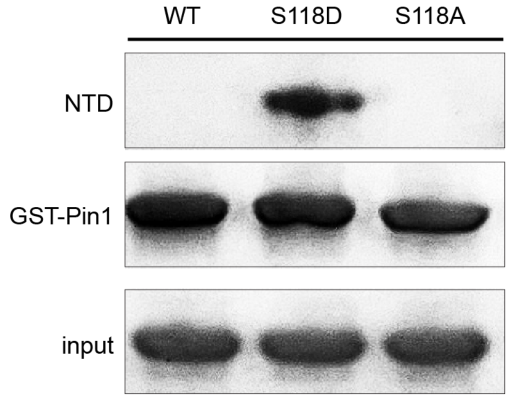


**Fig. S6. Validation of the phosphomimetic S118D mutation through GST-Pin1 pulldowns of purified proteins**.

GST-Pin1 pulldown assays were conducted, where the peptidyl-prolyl isomerase Pin1 was specially examined, given its known interaction with the Ser118-phosphorylated variant but not with the unphosphorylated wild-type or the phosphorylation-deficient S118A mutant[2]. SDS-PAGE analysis, stained with Coomassie blue, was consistent with previous findings and confirmed that Pin1 selectively interacts with the S118D mutant while demonstrating no interaction with the wild-type (WT) or the S118A variant.

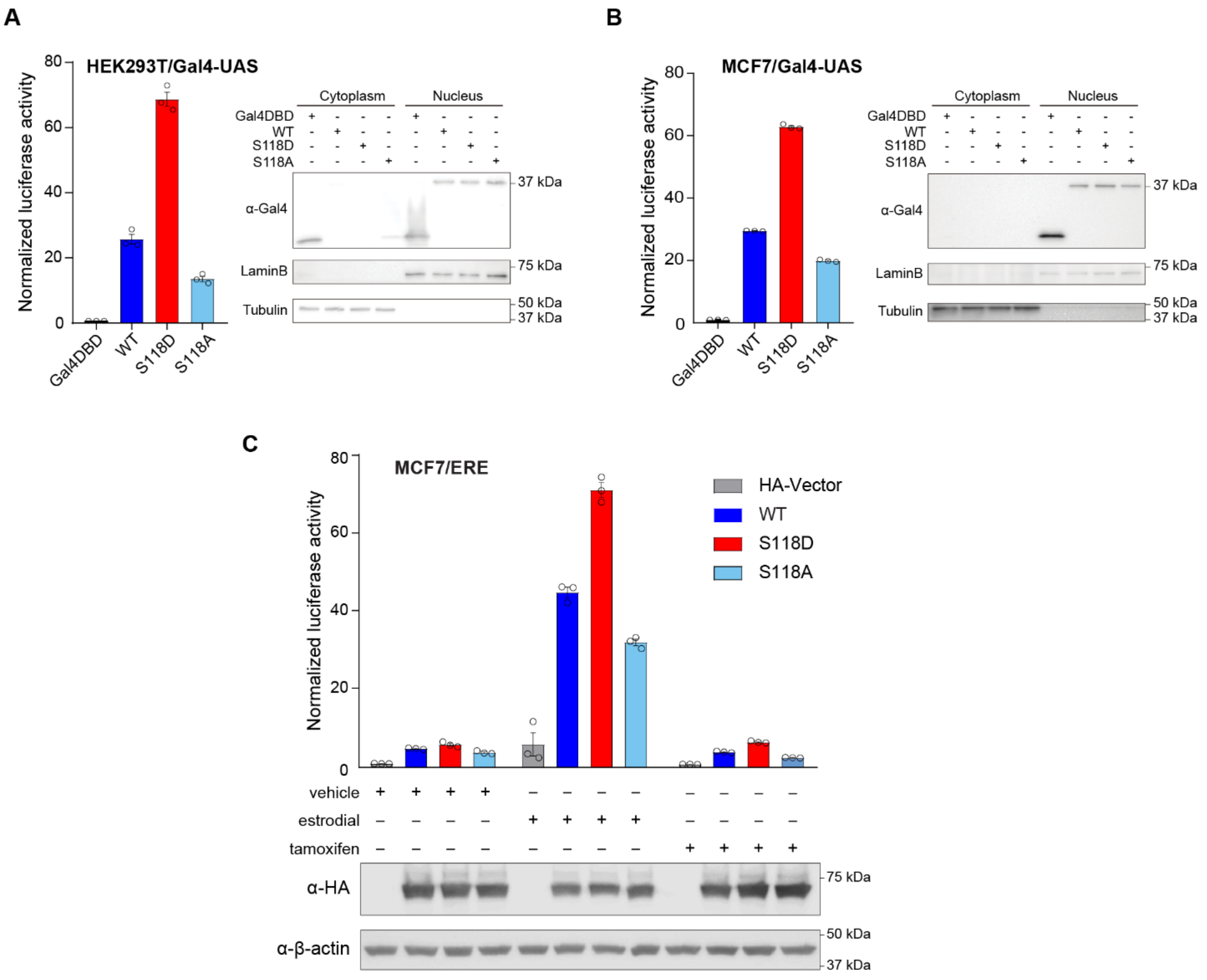


**Fig. S7. Activity enhancement by phosphomimetic S118D mutation for the NTD and full-length ER.**

(**A** and **B**) The transcriptional activity of Gal4-NTD was evaluated in HEK293T cells (A) and MCF7 cells (B) expressing wild-type (WT), S118D, and S118A variants. Right, western blot analysis of protein levels.

(**C**) The transcriptional activity of the full-length ER examined in MCF7 cells expressing wild-type (WT), S118D, and S118A variants. The cells were treated with ethanol (vehicle), 100 nM 17β-estradiol, and 1 μM tamoxifen for 12 hours. Mean ± SEM, calculated from 3 biological repeats. Bottom, western blot analysis of protein levels.

Transient transfections and dual-luciferase reporter assays were conducted to measure the transcriptional activity. The activity was determined by assessing Gal4 firefly luciferase levels, subsequently normalized by *Renilla* luciferase levels.

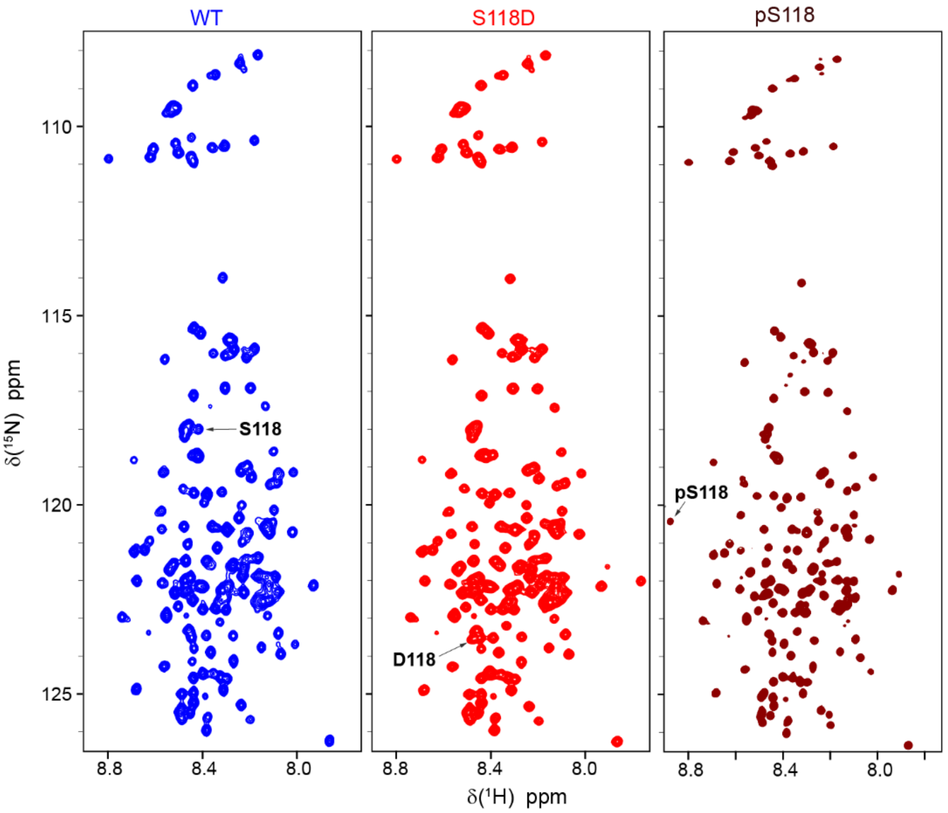


**Fig. S8. $^1$H-$^{15}$N HSQC spectra of wild-type ER-NTD and the S118D and pS118 variants.**

Deposited assignments for the wild-type ER-NTD are available in the Biomolecular NMR database.

For the S118D variant, deposited assignments are available in the Biomolecular NMR database.

For the pS118 variant, deposited assignments are available in the Biomolecular NMR database.

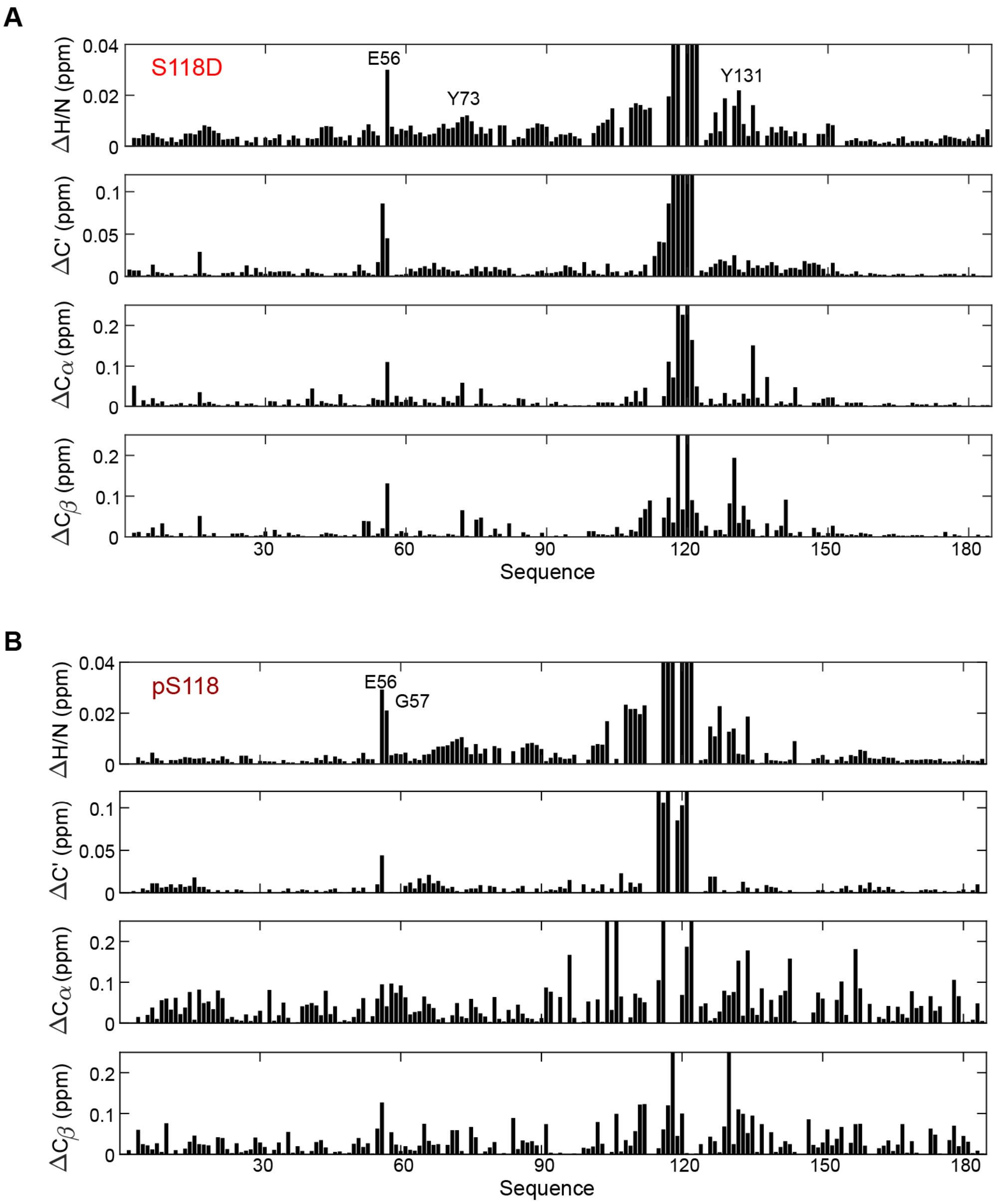


**Fig. S9. Long-range chemical shift changes induced by Ser118 phosphorylation and phosphomimetic S118D mutation.**

Chemical shift changes of ER-NTD amino acids were calculated using $^{1}H$, $^{15}N$, $^{13}C$ ($C^{\alpha}$, $C^{\beta}$, and $C'$) chemical shifts for the S118D variant (**A**) and the pS118 variant (**B**) relative to the wild-type. $\Delta H/N = \sqrt{(\Delta\delta(^{1}H))^2 + (\Delta\delta(^{1}N)\times 0.154)^2}$ , where Δδ is the chemical shift change for $^{1}H$ and $^{15}N$, respectively.

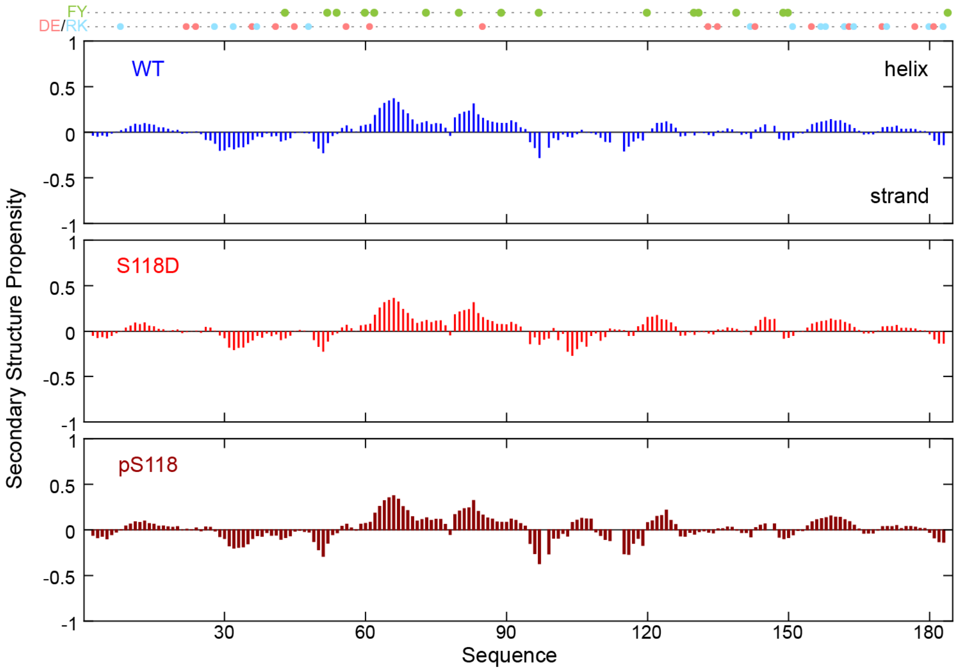


**Fig. S10. Lack of significant impact on secondary structure from Ser118 phosphorylation and phosphomimetic S118D mutation.**

Secondary structure propensity (SSP) was calculated for wild-type, S118D, and pS118, using their $^{1}H$, $^{15}N$, $^{13}C$ ($C^{\alpha}$, $C^{\beta}$, and $C'$) chemical shifts[3]. Low SSP scores indicate an absence of persistent secondary structure.

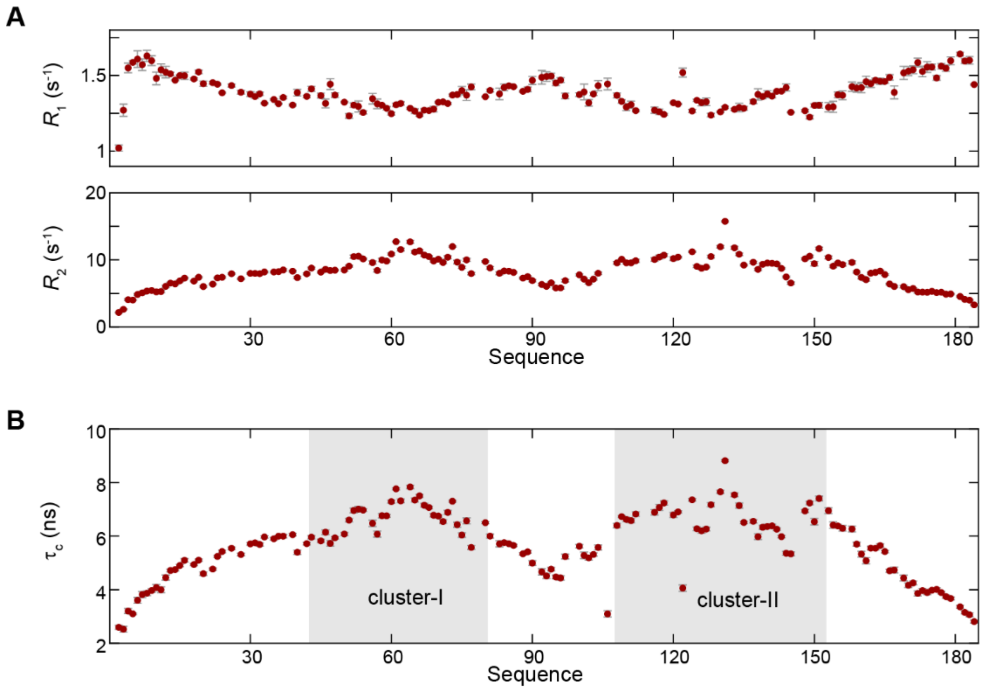


**Fig. S11. $^{15}$N relaxation measurements of the pS118 variant.**

**(A)** The profiles of $^{15}$N-$R_1$ and $^{15}$N-$R_2$ at 850 MHz for the pS118 variant.

**(B)** The residue-specific correlation time ($\tau_c$) determined from the measured $^{15}$N relaxation rates.

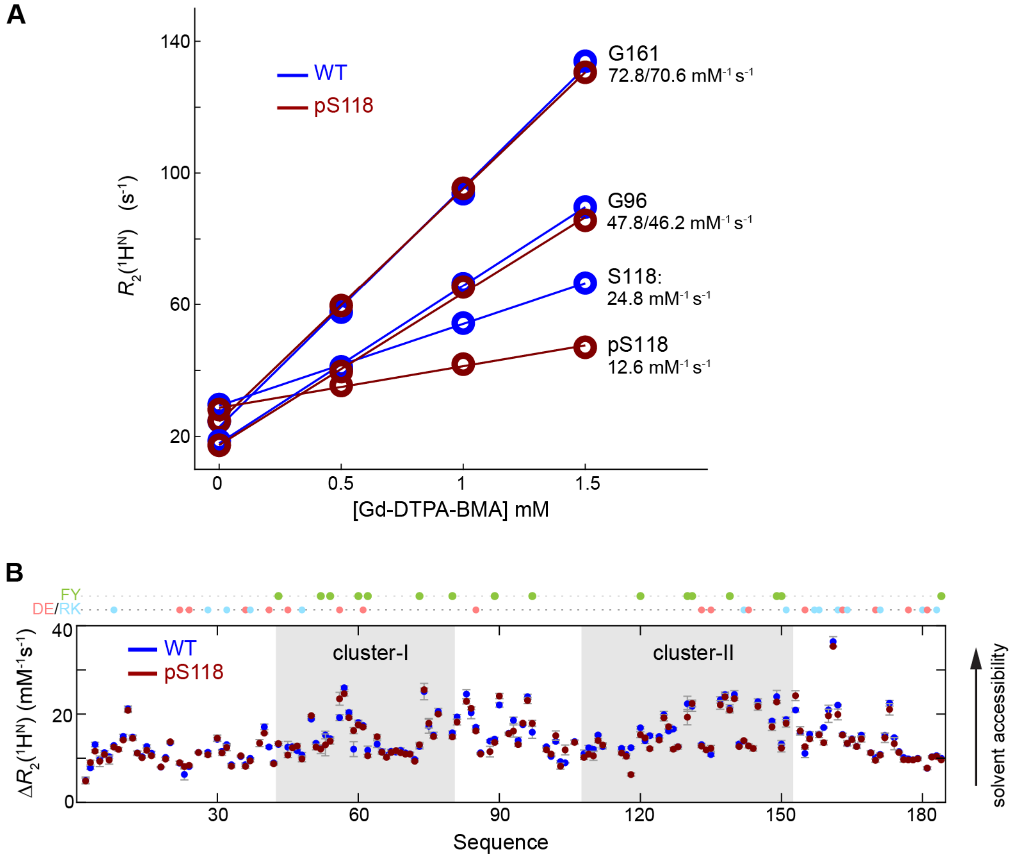


**Fig. S12. Solvent accessibility probed by soluble paramagnetic probes.**

(**A**) Freely diffusive paramagnetic Gd-DTPA-BMA probes were used to probe the residue-specific solvent accessibility. Transverse proton relaxation $R_2(^1H^N)$ rates were measured at a series of Gd-DTPA-BMA concentrations, as plotted for representative residues of both the wild-type (in blue) and the pS118 variant (dark red) at a protein concentration of 120 µM. The linear dose-response relationship provides information on the relative solvent accessibility of each residue, specifically the amide proton[4-6]. The slope of the linear relationship, expressed in units of $mM^{-1}$ $s^{-1}$, serves as a quantitative measure of solvent accessibility.

(**B**) Comparison of solvent accessibility (bottom) in the wild-type and the pSer118 variant. The solvent accessibility was derived using linear slope analysis of $R_2(^1H^N)$ at increasing concentrations of 0, 0.5, 1.0, and 1.5 mM of gadodiamide.

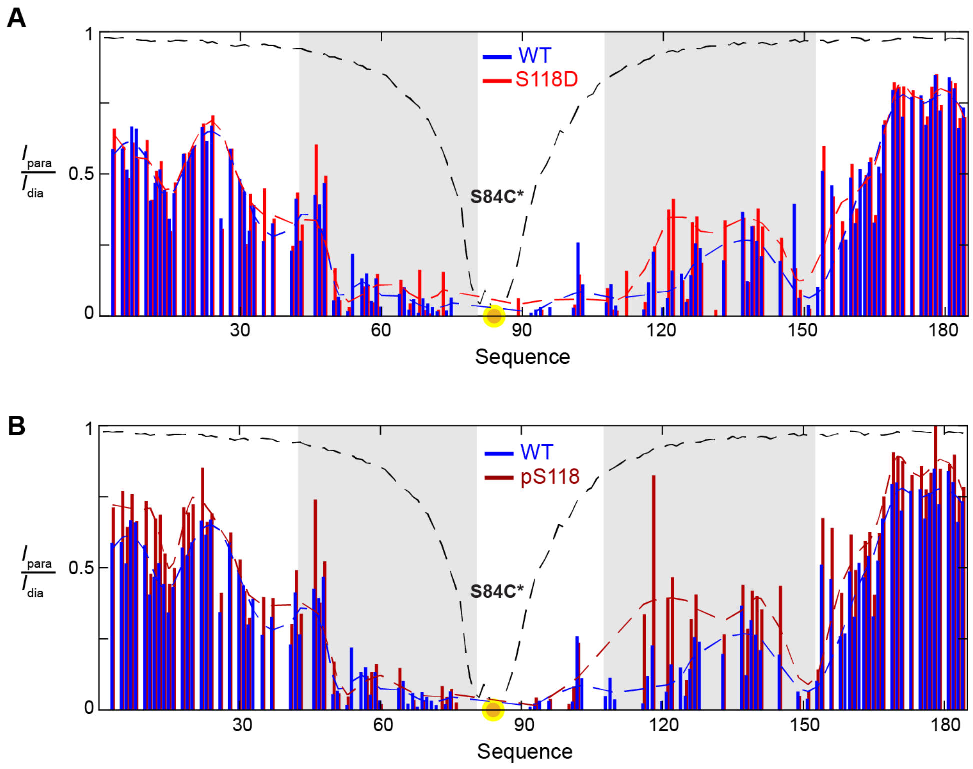


**Fig. S13. Phosphomimetic S118D mutation and S118 phosphorylation modulate long-range interactions probed by S84C spin labeling.**

**(A)** The PRE profiles of the wild-type (blue) and the S118D variant (red) are compared.

**(B)** The PRE profile of the pS118 variant (dark red) is compared to the wild-type (blue).

A protein concentration of 30 µM was used in both S118D and pS118 measurements. Yellow circle, the position of S84C spin labeling; dashes lines, visual aid. Yellow circle, the position of S84C spin labeling; dashes lines, visual aid.

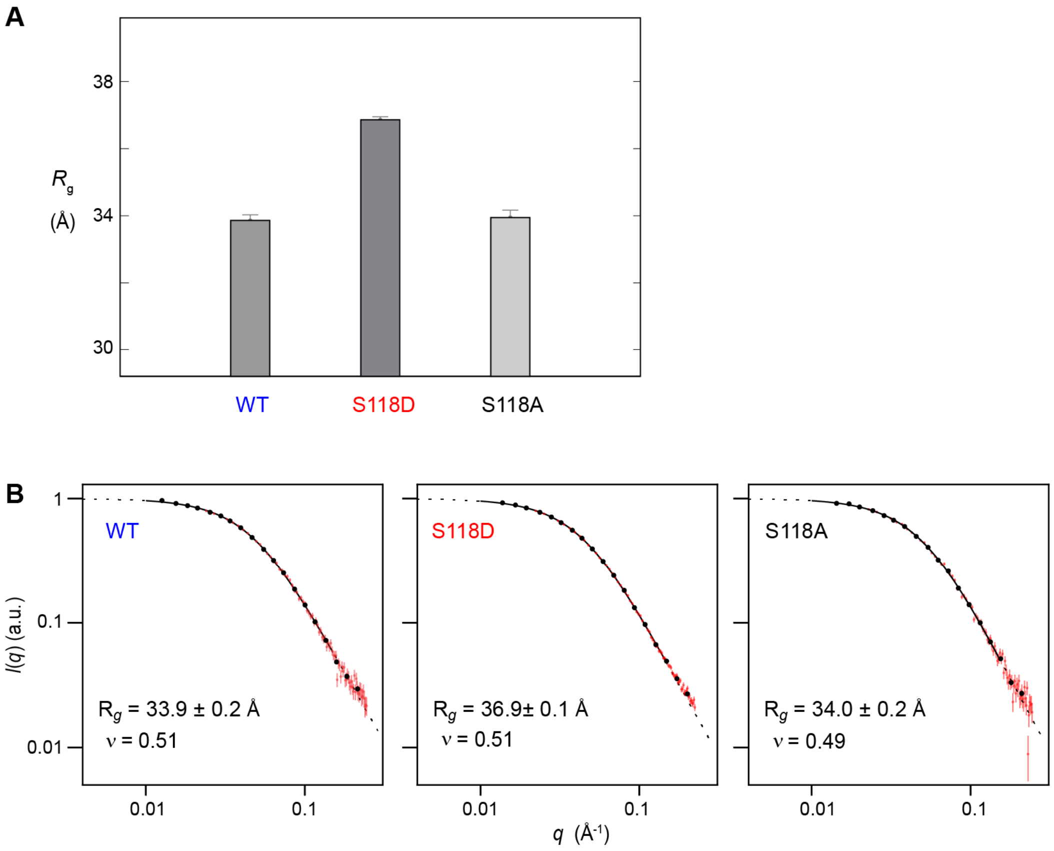


**Fig. S14. Phosphomimetic S118D mutation induces a 3-Å increase in radius of gyration.**

**(A)** Radius of gyration ($R_g$) of the wild-type (WT), the S118D, and the S118A variant, derived from SEC-SAXS data.

**(B)** Experimental SEC-SAXS profiles for the wild-type (left) and the S118D variant (center), as well as the S118A control (right), were acquired at 4°C using a pre-cooled running buffer for size-exclusion chromatography. $I(q)$, scattering intensity with error bars (normalized by zero-angle scattering); $q$, the amplitude of the X-ray scattering vector relating to the scattering angle. Dash lines, fitting results from the empirical molecular form factor (MFF) method[7], along with their resulting $R_g$ and $\nu$ values.

The SEC-SAXS data for the wild-type and the S118D and S118A variants are available in the Small Angle Scattering Biological Databank.

1. Huang, W., Peng, Y., Kiselar, J., Zhao, X., Albaqami, A., Mendez, D., Chen, Y., Chakravarthy, S., Gupta, S., Ralston, C., Kao, H.-Y., Chance, M.R. & Yang, S. Multidomain architecture of estrogen receptor reveals interfacial cross-talk between its DNA-binding and ligand-binding domains. *Nature communications* **9**, 3520 (2018).

2. Rajbhandari, P., Finn, G., Solodin, N.M., Singarapu, K.K., Sahu, S.C., Markley, J.L., Kadunc, K.J., Ellison-Zelski, S.J., Kariagina, A., Haslam, S.Z., Lu, K.P. & Alarid, E.T. Regulation of estrogen receptor alpha n-terminus conformation and function by peptidyl prolyl isomerase pin1. *Mol Cell Biol* **32**, 445-457 (2012).

3. Tamiola, K., Acar, B. & Mulder, F.A. Sequence-specific random coil chemical shifts of intrinsically disordered proteins. *J Am Chem Soc* **132**, 18000-18003 (2010).

4. Gong, Z., Schwieters, C.D. & Tang, C. Theory and practice of using solvent paramagnetic relaxation enhancement to characterize protein conformational dynamics. *Methods* **148**, 48-56 (2018).

5. Hocking, H.G., Zangger, K. & Madl, T. Studying the structure and dynamics of biomolecules by using soluble paramagnetic probes. *Chemphyschem : a European journal of chemical physics and physical chemistry* **14**, 3082-3094 (2013).

6. Kooshapur, H., Schwieters, C.D. & Tjandra, N. Conformational ensemble of disordered proteins probed by solvent paramagnetic relaxation enhancement (spre). *Angew Chem Int Ed Engl* **57**, 13519-13522 (2018).

7. Riback, J.A., Bowman, M.A., Zmyslowski, A.M., Knoverek, C.R., Jumper, J.M., Hinshaw, J.R., Kaye, E.B., Freed, K.F., Clark, P.L. & Sosnick, T.R. Innovative scattering analysis shows that hydrophobic disordered proteins are expanded in water. *Science* **358**, 238-241 (2017).

8. Svergun, D.I. Determination of the regularization parameter in indirect-transform methods using perceptual criteria. *J Appl Crystallogr* **25**, 495-503 (1992).
